# Rainfall immediately before and after fire promotes long-term occurrence of a rare, fire-sensitive passerine

**DOI:** 10.64898/2026.03.03.709440

**Authors:** William F. Mitchell, David Paton, Rohan H. Clarke, Jemima Connell, Simon Verdon

## Abstract

Attributes of fire regimes are known to drive habitat suitability for many species in fire-prone environments. Comparatively little is known about how abiotic conditions (e.g. rainfall events, cumulative rainfall, drought) at the time of fire may affect long-term (>2-years) post-fire occurrence. We sought to a) establish whether the post-fire development of heathland habitat for the endangered mallee emu-wren is influenced by rainfall within 12-months before or after the most-recent fire, b) identify the preferred fire-age of heathland vegetation for the mallee emu-wren, and c) map those habitats most likely to support the species across a large reserve (∼271,000 ha), Ngarkat Conservation Park, from which it has been extirpated.

Using historical presence records, collected prior to the extirpation of mallee emu-wrens from the study area, we implemented a random-forest modelling approach to predict relative likelihood of occurrence (considered a proxy for probability of suitable habitat).

Rainfall in the 12-months before and after fire had a positive effect on relative likelihood of mallee emu-wren occurrence. The development of high-quality mallee emu-wren habitat required at least 420 mm of rainfall in the 12-months prior to the most recent fire. Only 35% of Ngarkat received rainfall above this threshold prior to the most recent fire. Rainfall in the 12-months after fire positively influenced relative likelihood of mallee emu-wren occurrence, though the effect was less pronounced than pre-fire rainfall. Relative likelihood of mallee emu-wren occurrence peaked 15 years after fire, with an ∼10-year peak time window of relative occurrence (10-20-years).

This study highlights that abiotic conditions at the time of fire, particularly rainfall in the 12-months preceding fire, have long-lasting impacts on relative probability of occurrence for this fire-sensitive species. Targeting fire management in ways that maximise post-fire occurrence of the mallee emu-wren – particularly by burning senesced habitat following periods of elevated rainfall – has potential to enhance conservation outcomes. Given the substantial and long-term impact of rainfall around the time of a fire identified in this study, short-term climatic conditions deserve greater attention in a range of ecosystems where managers aim to use fire to manipulate habitat for the benefit of fire-sensitive species.

## Introduction

Fire influences flora and fauna composition in many ecosystems globally, through complex mechanisms that depend on the traits of fire and the ecological context in which a fire occurs (Bowman et al. 2009, Gibson et al. 2013, He et al. 2019). The term ‘fire regime’ is used to capture the combination of fire characteristics of a given area (e.g., fire severity, frequency, size and season), with frequent focus on those attributes related local ecosystem dynamics (Wright and Clarke 2007, Krebs et al. 2010, Archibald et al. 2013, Harvey et al. 2016). Time since fire has been afforded considerable research attention, due to its fundamental role as an abrupt disturbance event which resets the process of vegetation succession, thereby driving habitat suitability for many species in fire-prone environments (Smucker et al. 2005, Kenny et al. 2018, Clarke et al. 2021).

Biotic and abiotic factors such as climate, local weather events, landscape characteristics and inter-taxa relationships also influence species responses to fire (Gibson et al. 2013, Giljohann et al. 2017, Kenny et al. 2018, Connell et al. 2022). Abiotic conditions influence biotic responses to fires directly, while also interacting with fire regime components to shape species responses to fire. Despite the ecological significance of abiotic conditions such as rainfall at the time of a fire, these factors have received little research attention. This may be because such short-duration events occur unpredictably and with highly variable intensity, making them difficult to incorporate into planning and analysis.

Interactions between fire and discrete abiotic conditions have the potential for considerable and long-term impacts on ecosystem structure. For example, the compounding effects of large fire and drought within a short time period have the potential to trigger sudden shifts in vegetation community state (Batllori et al. 2018). For habitat specialists, such a transition may represent the loss of considerable areas of once-favourable habitat. Weather conditions are an important determinant of regeneration rates in the years immediately following fire (Giljohann et al. 2017, Dikshit and Evans 2024, Volkova et al. 2025). The same can be true prior to fire. In an Australian arid fire-prone system, post-fire recruitment of the obligate-seeding grass soft spinifex *Triodia pungens* is strongly associated with elevated seed bank density resulting from above-average rainfall prior to fire (Wright and Fensham 2018). Nevertheless, it is often unclear whether the short-term effects of weather conditions on post-fire regeneration confer a long-term influence on the condition of ecological communities, or their potential as habitat for other species.

Understanding how abiotic conditions influence the ecological impact of fire is increasingly important due to climate change (Jones et al. 2022, Grau-Andrés et al. 2024, Charles et al. 2025). The frequency of drought and extreme weather events are increasing in many parts of the world, including many fire-prone ecosystems, while other areas are predicted to receive increased rainfall and milder weather (Hope et al. 2015). Regardless of direction, climatic shifts will influence both the probability and severity of fire, and the abiotic conditions at the time of fires (Canadell et al. 2021).

Managing fire to maintain or create habitat may be a critical step in the conservation of taxa with limited capacity to adapt to the compounding impacts of altered fire regime, climate change, and habitat destruction (Puig-Girones et al. 2025). Fire has been used as a management tool for many thousands of years, and is supported by an extensive and global body of research (Driscoll et al. 2010), with a frequent focus on identifying fire-history traits that maximise biodiversity or viability of threatened taxa at a landscape scale (e.g., Kelly et al. 2015, Verdon and Clarke 2022, Puig-Girones et al. 2025). Many such studies incorporate abiotic conditions at the time of sampling (e.g., Arthur et al. 2012, Connell et al. 2022) or consider the influence of abiotic conditions on short-term post fire recovery of taxa (e.g., Yarnell et al. 2007, Plavsic 2014). However, it is unclear whether discrete abiotic events within a relatively short time of fire have a long-term influence on habitat condition. The ecological consequences of decisions to burn or to suppress fire may persist for decades or even centuries under long inter-fire intervals (Connell et al. 2017). For species which require long unburnt habitat (e.g., the black-eared miner *Manorina melanotis*; Clarke et al. 2005), such consequences may not be apparent until decades after management decisions are made (Connell et al. 2017). With fire season length predicted to increase under climate change in many parts of the world (Jones et al. 2022), the importance of adopting fire management strategies that most effectively promote habitat and populations of fire-sensitive species becomes ever greater.

To test the potential impacts of abiotic conditions at the time of fire on long-term post-fire recovery, we studied the mallee emu-wren *Stipiturus mallee* in a fire-prone region in south-eastern Australia. The mallee emu-wren is an endangered, fire and drought sensitive passerine with a narrow geographic range and ecological characteristics that make it vulnerable to changes in vegetation composition associated with altered fire regimes (Indigo et al. 2023). Using historical records, collected prior to the extirpation of the species from much of its remnant heathland habitat, we sought to establish whether the development of appropriate heathland habitat for this species was dependent on cumulative rainfall within the 12-month period before or after the most recent fire. Additionally, we sought to determine the preferred fire-age for this species in heathlands and to map the distribution of potential mallee emu-wren habitat across the 271,000-ha study area.

## Methods

### Study system and focal species

We focused this study in a vast (271,000 ha) area of contiguous native vegetation characterised by fire-prone Mediterranean shrublands (Figure 1). The area is protected as Ngarkat Conservation Park (Ngarkat) and predominantly supports two broadly defined vegetation groups: heathlands and mallee shrublands or woodlands (Australian Government Department of Climate Change Energy the Environment and Water 2023). Within this landscape, fine scale variability in vegetation structure and composition is driven by topography, particularly dune-swale systems, soil geology, fire regime, and a regional climate gradient. The southwest receives the greatest rainfall with an annual median of 451 mm and median maximum temperature of 22.3 °C (Keith weather station, Bureau of Meteorology (BOM) 2025), while the driest areas in the northeast receive median annual rainfall of 327 mm (Pinnaroo weather station; BOM 2025) and a median maximum temperature of 23.5 °C (Lameroo weather station, BOM 2025). Characteristic of Mediterranean shrublands the world over, this system experiences hot dry summers and relatively frequent fire, with many inhabitant plant species exhibiting fire-tolerant or fire-dependent adaptations (Department for Environment and Heritage 2009). Summer thunderstorms are not uncommon and are the dominant cause of wildfire (Keith et al. 2002, Department for Environment and Heritage 2009).

**Figure 1.**
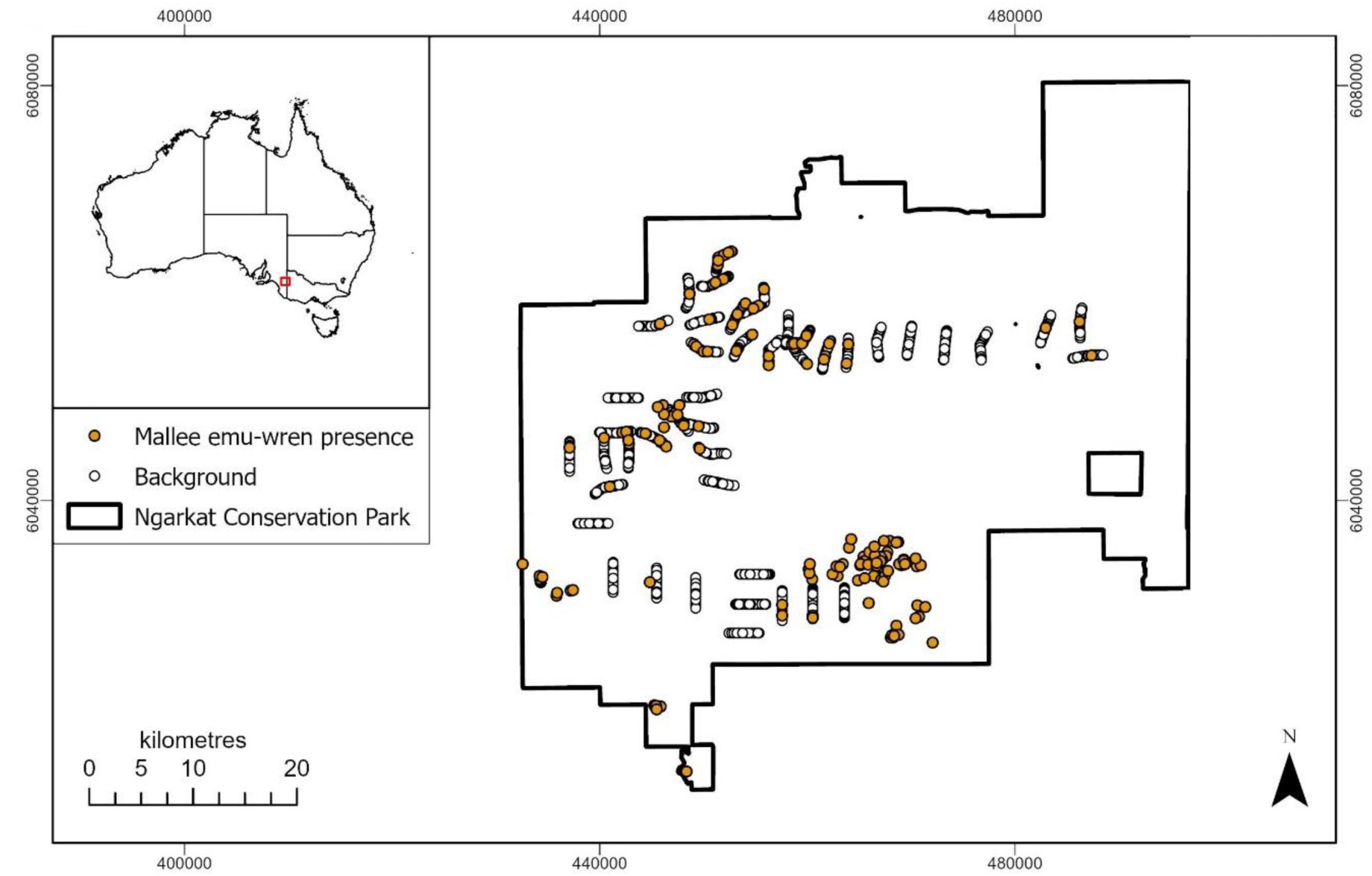
Location of presence and background records used in this study. Inset, study region location within Australia.

Many locations within this study region once supported mallee emu-wrens (Figure 1). The mallee emu-wren typically inhabits two distinct vegetation types: Triodia mallee (a subset of mallee shrublands and woodlands characterised by open multi-stemmed Eucalypts to ∼7 m with an understorey dominated by the hummock grass *T. scariosa*) and heathlands (characterised by few or no trees, dense cover of xenomorphic shrubs and, compared with Triodia mallee, reduced density of *Triodia scariosa*). In either vegetation type, the mallee emu-wren has a strong association with *T. scariosa* (Verdon et al. 2020) which has a patchy distribution in this landscape. In Triodia mallee vegetation, the distribution of mallee emu-wren habitat is spatially and temporally dynamic, with fire history, rainfall and local elevation driving variability in habitat suitability over time (Verdon et al. 2019, Connell et al. 2022). After fire, *T. scariosa* requires ∼15 years to attain sufficient structure to support mallee emu-wrens (Brown et al. 2009). After 20 – 30 years *T. scariosa* size and cover will plateau or gradually decline (Haslem et al. 2011). Without fire to reset the process of succession, *T. scariosa* may senesce to the point where it no longer supports mallee emu-wrens. Successive large wildfires in the 21^st^ century have led to the species’ extirpation from Ngarkat, though a warming climate and habitat fragmentation have also contributed to declines across the global mallee emu-wren population (Indigo et al. 2023). In 2025, the mallee emu-wren is restricted to only three spatially disconnected conservation reserves, all in south-eastern Australia. Within heathland vegetation, only a single small, isolated population is known to persist, ∼60 km east of Ngarkat (Verdon et al. 2025). Consequentially, little is known about the species’ fine-scale habitat requirements or fire response in heathlands when compared with Triodia mallee. In the twenty years preceding this study, 80% of Ngarkat has been burnt at least once, including the last area known to support the mallee emu-wren in 2014. The mallee emu-wren is the current focus of coordinated and large-scale conservation efforts by multiple organisations, including *in-situ* conservation initiatives and a long-term plan to reintroduce the species into areas of its historical range which have regenerated following fire (Mitchell et al. 2021, Indigo et al. 2023).

### Mallee emu-wren presence records

We pooled mallee emu-wren presence records from multiple survey programs, as well as incidental records, from the period 1990-2014 (data provided by D Paton and R Clarke and constituted transect surveys, as well as incidental records of mallee emu-wrens ‘off transect’). We also pooled presence records of other bird species from the same period to use as ‘background’ records (background records were collected up to 2016 to allow comparison with Big Desert, see supplementary material). Background data points were restricted to records made during surveys that would have recorded mallee emu-wrens if present (i.e. surveys of all bird species or of threatened bird species). Because mallee emu-wrens are cryptic and small, targeted surveys typically include the broadcast of pre-recorded mallee emu-wren vocalisations to elicit a response and increase detectability (Mitchell et al. 2021, Verdon et al. 2025). Vocalisation broadcasting was not routinely employed during data collection for this study and for this reason, we did not consider ‘background’ points as true absences. In cases of multiple records within 100 m within a three-year period, we avoided duplicates by retaining only a single record. A comprehensive history of fires between 1940 and 2020 for the region was sourced from the South Australian Government’s ‘Bushfires and Prescribed Burns History’ dataset (Department of Environment and Water 2020). Records of fires prior to 1940 were not available for the study region. For this reason, datapoints where the most recent fire occurred before 1941 were filtered from the dataset (4 presence, 269 ‘background’ records). For a small subset of fires, particularly in the mid-20^th^ century, only the year, and not the date of fire was available. This resulted in uncertainty about the most recent fire for a small number of data points. All such examples were removed (1 presence, 42 ‘background’ records). The final dataset included 208 mallee emu-wren records and 15,087 ‘background’ records.

### Predictor variables expected to influence relative likelihood of mallee emu-wren occurrence

We considered several candidate variables expected to influence mallee emu-wren occurrence (variables associated with fire, vegetation cover, topography, climate and local geology, table 1) and prepared raster layers for each variable covering the extent of the study region with a pixel size of 75 m x 75 m (the finest resolution available for all candidate variables) (ESRI 2024).

**Table 1.** variables considered as potential predictors of mallee emu-wren occurrence in Ngarkat Conservation Park, South Australia.

| Covariate Name | Description | Rational for inclusion | Data source | Included in final model? |
| --- | --- | --- | --- | --- |
| Years since fire | Number of calendar years since most recent fire at the time of species record. | Known to influence Mallee emu-wren occurrence (Brown et al. 2009, Verdon et al. 2019). | Bushfires and Prescribed Burns History South Australia (Department of Environment and Water 2020) | Yes |
| Hectares of heathland vegetation in surrounding area | Total number of 75 x 75 m raster cells categorised as 'heathland' within a radius of 12 cells (~900 m). | A landscape-scale measure of extent of potential habitat (Paton et al. 2009, Haslem et al. 2010, Verdon et al. 2025). | National Vegetation information System (Australian Government Department of Climate Change Energy the Environment and Water 2023) | No (low importance) |
| Localised elevation (height above rivers) | An index of elevation where the baseline is set by the mean height of rivers in the region rather than metres above sea level. | Local elevation can influence mallee emu-wren habitat suitability (Verdon et al. 2019). | Digital elevation model (Gallant et al. 2009). | Yes |
| Hill shade | Combines aspect and slope to calculate exposure to sun under a given set of conditions. For calculations, we located the sun at 45 degrees above the horizon and north (0 degrees). | Expected to affect microclimate and composition of habitat-forming vegetation species | Digital Elevation Model (Gallant et al. 2009). | No (low importance) |
| Wind exposure index | An index representing local topographic position (from exposed dune-top to sheltered swale). Based on the elevation and aspect of a cell compared to the surrounding eight raster cells. Values below 1 indicate wind shadowed areas whereas values above 1 indicate areas exposed to wind (Böhner and Antonić 2009, Gerlitz et al. 2015). | Expected to affect microclimate and composition of habitat-forming vegetation species | Elevation Model (Gallant et al. 2009). | No (low importance) |
| Terrain ruggedness | An index of topographic complexity representing the elevational difference between each location (raster cell) and the surrounding terrain (raster cells). Ruggedness at each location was assessed by comparing it to the surrounding eight raster cells. | Expected to affect microclimate and composition of habitat-forming vegetation species | Digital Elevation Model (Gallant et al. 2009) | No (low importance) |
| Diurnal Anisotropic heat | Estimate of anisotropic heat distribution: $\text{Anisotropic heat} = \cos(\text{amax} - a) * \arctan(b)$ where amax = aspect with the maximum total heat surplus, a = aspect and b = slope (Böhner and AntoniĆ 2009). | Expected to affect microclimate and composition of habitat-forming vegetation species | Digital Elevation Model (Gallant et al. 2009). | No (low importance) |
| Mean annual rainfall | National, gridded mean annual rainfall is calculated over 30 years from weather station data, satellite data, and modelling. | Represents the climate gradient across the study area, with mallee emu-wrens expected to be more resilient in cooler, wetter areas. | Bureau of Meteorology (http://www.bom.gov.au). | Yes |
| Aspect | The compass direction a slope is facing. Averaged over each raster cell. | Expected to affect microclimate and composition of habitat-forming vegetation species | Digital Elevation Model (Gallant et al. 2009) | Yes |
| Rainfall in the 12 months prior to most recent fire | Cumulative rainfall in the 12-months prior to most recent fire at the time of species record. |  | Keith weather station (Bureau of Meteorology 2024) | Yes |
| Rainfall in the 12 months following most recent fire | Cumulative rainfall in the 12-months following most recent fire at the time of species record. |  | Keith weather station (Bureau of Meteorology 2024) | Yes |
| Thorium content | Remote-sensed radiometric measurement correlated with soil clay content in the study area. | A proxy for clay content in soil. Soil composition expected to affect composition of habitat-forming vegetation species (Verdon et al. 2019). | (Read et al. 2018) | Yes |
| Vegetation group | National Vegetation Information System<br>Major Vegetation Group layer | Known to influence Mallee emu-wren occurrence (Brown et al. 2009, Verdon et al. 2019). | National Vegetation information System (Australian Government Department of Climate Change Energy the Environment and Water 2023). | No (low importance) |
| Clay (0 – 5 cm) | Clay content of soil at depth of 0 – 5 cm | Soil composition expected to affect composition of habitat-forming vegetation species (Verdon et al. 2019). | Soil and Landscape Grid National Soil Attribute Map – Clay (Malone and Searle 2022) | No (low importance) |
| Clay (5 – 15 cm) | Clay content of soil at depth of 5 – 15 cm | See above | See above | No (correlation) |
| Clay (15 – 30 cm) | Clay content of soil at depth of 15 – 30 cm | See above | See above | No (correlation) |
| Clay (30 – 60 cm) | Clay content of soil at depth of 30 – 60 cm | See above | See above | No (correlation) |
| Clay (60 – 100 cm) | Clay content of soil at depth of 60 – 100 cm | See above | See above | No, correlation |
| Clay (100-200 cm) | Clay content of soil at depth of 1 – 2 m | See above | See above | No, low importance |

To calculate a temporally accurate value for each of the three variables associated with fire included in this study (years since fire, cumulative rainfall in the 12-months prior to most recent fire and cumulative rainfall in the 12-months following most-recent fire) we first determined the month and year of the most recent fire at the time that each datapoint was collected. Rainfall data for these variables was then obtained from the Bureau of Meteorology weather station located in the town of Keith, approximately 15 km from the southwestern boundary of the study region (Bureau of Meteorology 2024). We tested for correlations between all candidate variables and excluded those that were highly correlated (> 0.70). For some fires, particularly in the mid-20^th^ century, whether a fire was a planned burn or wildfire was not known and thus whether a fire was wild or controlled, or any measure of fire severity, were unavailable for inclusion in our analysis, despite the importance of these aspects of fire regime (e.g., Smucker et al. 2005, Doherty et al. 2024).

### Analyses

*Random forests* is a machine learning approach with broad uptake as a method for developing species distribution models (SDMs, Valavi et al. 2021). Random forest models are non-parametric, inherently capture interactions between predictors, and allow the importance of predictors to be estimated (Valavi et al. 2021). These features make it a flexible and powerful tool when working with ‘presence-only’ data that has been collected in a ‘semi-structured’ manner, as is the case in this study. Random forests with down sampling has been shown to be amongst the top performing commonly used approaches for species distribution modelling (Valavi et al. 2022). The use of historical records to make predictions about contemporary habitat suitability or species occurrence may be misleading if the environmental conditions at the time records were made were associated with a population in decline, or if environmental conditions have changed in the period between data collection and contemporary modelling (Bracken et al. 2022). However, training species distribution models using historical presence records and environmental variables with high persistency over relevant time scales (e.g., geographical or geological data) can yield high predictive performance (Bracken et al. 2022).

We used the package *randomForest* version 4.7-1.2 (Liaw and Wiener 2002) in the statistical environment *R* (version 4.4.2, R Core Team 2024) to generate a random forest classification model estimating relative likelihood of mallee emu-wren occurrence across our study region. This model was trained using presence/background data and as such, the response represents a relative likelihood of occurrence, rather than an absolute probability of occurrence (Guillera-Arroita et al. 2015). To account for bias caused by an imbalance between the number of presence and background records, we incorporated a down-sampling procedure in our modelling approach (i.e., each individual tree was fitted with a random sample containing the same number of presence and background points). Our first iteration random forest classifier included all candidate covariates which had not been excluded due to correlation and used 10,000 decision trees. Predictor variables were ranked by importance using ‘mean decrease in accuracy’ and ‘mean decrease in Gini impurity’. The *randomForest* package calculates ‘mean decrease in accuracy’ by randomly permuting each predictor in the out of bag (OOB) data and measuring how much the overall model accuracy declines (Liaw and Wiener 2002). Higher values indicate a greater contribution to model performance for a given predictor. OOB data refers to the subset of training data that is not included in the bootstrap sample for a given tree and can therefore be used to validate model predictions. Gini impurity quantifies how homogeneous the data are within a node of a decision tree (Liaw and Wiener 2002). Lower values indicate ‘purer’ nodes, where most observations belong to the same class. We retained only those variables ranked in the top 5 for either ‘mean decrease in accuracy’ or ‘mean decrease in Gini impurity’ and retrained the model. We compared the first iteration and improved random forest model using (OOB) error rate. This process was repeated iteratively with successively fewer variables retained, until model performance stopped improving. The final model yielded greater balanced accuracy (mean of presence and background error rates) than the first iteration classifier by ∼5%.

Ngarkat forms the western part of a broader expanse of contiguous native vegetation (∼980,000 ha) incorporating Big Desert Wilderness Park, Big Desert State Forest and Wyperfeld National Park (hereafter, these contiguous reserves, excluding Ngarkat, are referred to as Big Desert). Compared with Ngarkat, survey effort, and consequently mallee emu-wren records, are scarce in Big Desert. To assess the feasibility of extrapolating our model predictions across this broader region, we predicted relative likelihood of mallee emu-wren occurrence at each of 298 available data points (25 presence and 273 background records) and compared model predictions against mallee emu-wren presence/background data. As an additional validation test, we compared random forest predicted relative likelihood of mallee emu-wren occurrence between Ngarkat and Big Desert at 229 unique 25 ha sites which had been assigned a qualitative category regarding the presence of mallee emu-wren habitat by an experienced ecologist. For these tests, fire history data for Big Desert was sourced from the Victorian Government’s ‘Fire History Records of Fires across Victoria’ dataset (Department of Energy Environment and Climate Action 2023). In both tests, random forest model performance was considered poor across Big Desert when compared with Ngarkat, possibly due to the paucity of records in Big Desert, or ecological differences between the regions. Because of this poor performance, we made no attempt to extrapolate model predictions beyond Ngarkat (supplementary material).

Using the package *iml* (Molnar and Bischl 2018), we generated partial dependence (PD) and individual conditional expectation (ICE) plots to explore the effect of individual predictor variables associated with the most recent fire at any given location on relative likelihood of mallee emu-wren occurrence. PD shows the effect of a predictor on the modelled response, averaging over the observed distributions of all other predictors. ICE plots provide additional nuance by showing the effect of a predictor on the modelled response for each observation, while holding all other predictors at their observed values. To increase interpretability, we limited the plotted cases for all ICE plots to a random sample of 500. We also generated two-way PD plots to examine the interactions between years since fire and cumulative rainfall in the 12-months preceding and following fire using the *pdp* package (Greenwell 2017). Finally, we used the *raster* package (Hijmans 2025) in R and the most up-to-date fire history data for South Australia (Department of Environment and Water 2025), to project the predicted relative likelihood of mallee emu-wren occurrence across the 271,000 ha Ngarkat in December 2025. For two fires, that burnt in March and May 2025, we used median rainfall values for December 2025 and later months to calculate cumulative rainfall in the 12-months following fire. Because mallee emu-wrens have almost certainly been extirpated from Ngarkat (Verdon et al. 2025), we considered modelled relative likelihood of mallee emu-wren occurrence as a proxy for probability of suitable mallee emu-wren habitat, rather than representing the actual likelihood of birds being present. We also predicted relative likelihood of mallee emu-wren occurrence for the years 2035 and 2045, assuming no future fire.

## Results

Our final model performed well, predicting mallee emu-wren presences with an OOB error rate of 0.16 and background points with an OOB error rate of 0.11 (0 = 0% error, 1 = 100% error). Mean annual rainfall was positively associated with relative likelihood of mallee emu-wren occurrence (Figure 2). Greater rainfall in the 12-months both preceding and following fire had a positive effect on relative likelihood of mallee emu-wren occurrence (Figures 2 – 3). Mallee emu-wrens were most likely to occur 10-20 years post-fire where the total cumulative rainfall exceeded ∼420 mm in the 12 months prior to the most recent fire (Figures 2 – 3). Despite median annual rainfall of 451 mm (Keith weather station), only 35% of Ngarkat received cumulative rainfall above the 420 mm threshold in the 12-months prior to most recent fire. Cumulative rainfall exceeded this threshold in 41 of the 69 unique fire events which make up the current distribution of most-recent fires across Ngarkat. Relative likelihood of mallee emu-wren occurrence peaked 15 years after fire, with an ∼10-year time-window of occurrence (Figures 2 – 3).

**Figure 2.**
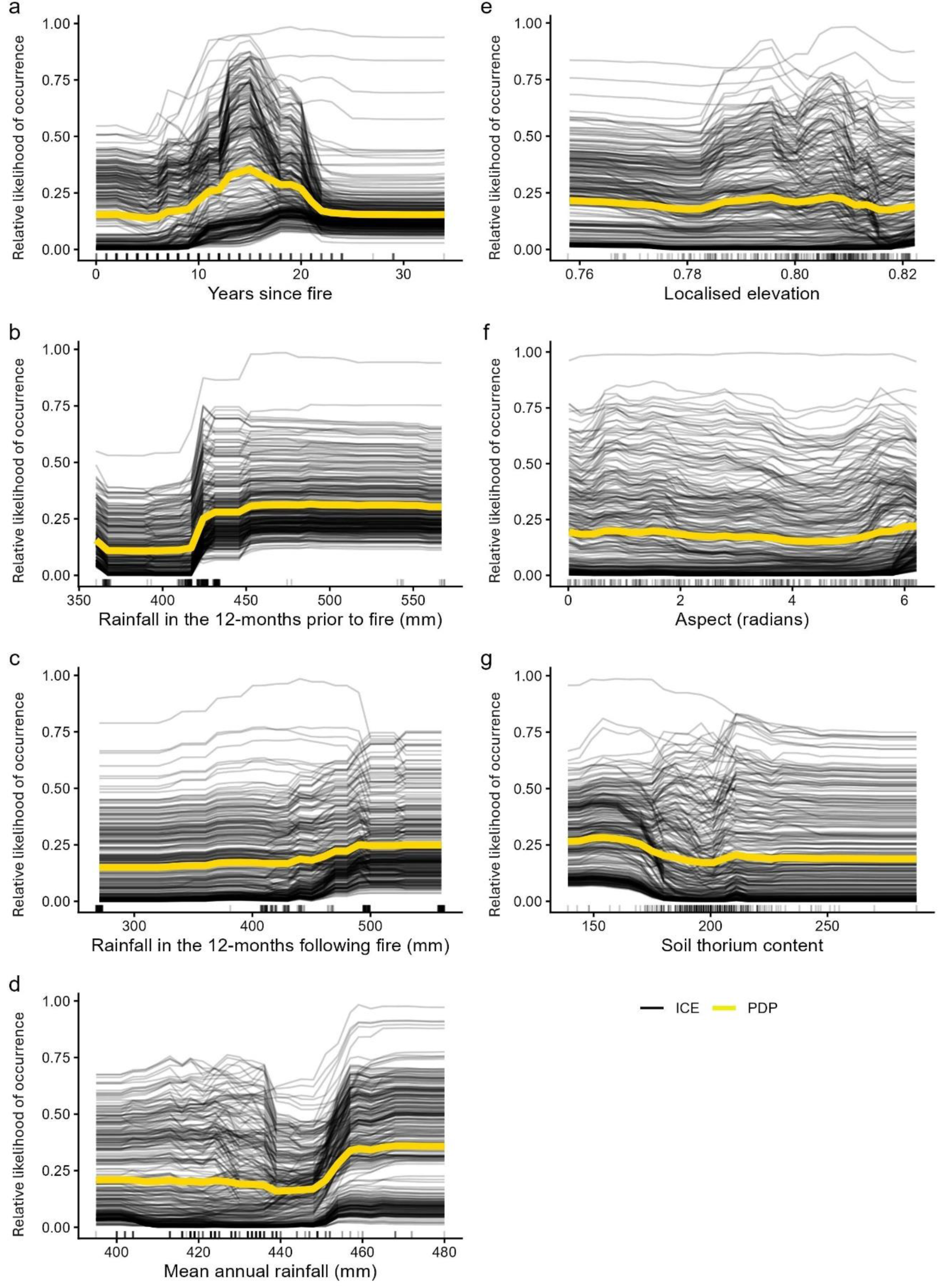
Individual conditional expectation (ICE) plot with partial dependence (PD) predicted relative likelihood of mallee emu-wren occurrence as a response to a) years since fire, b) rainfall in the 12-months prior to most recent fire and c) rainfall in the 12-months following most recent fire. To aid clarity, a random sample of 500 instances are shown for each ICE plot. Rug depicts individual data points.

**Figure 3.**
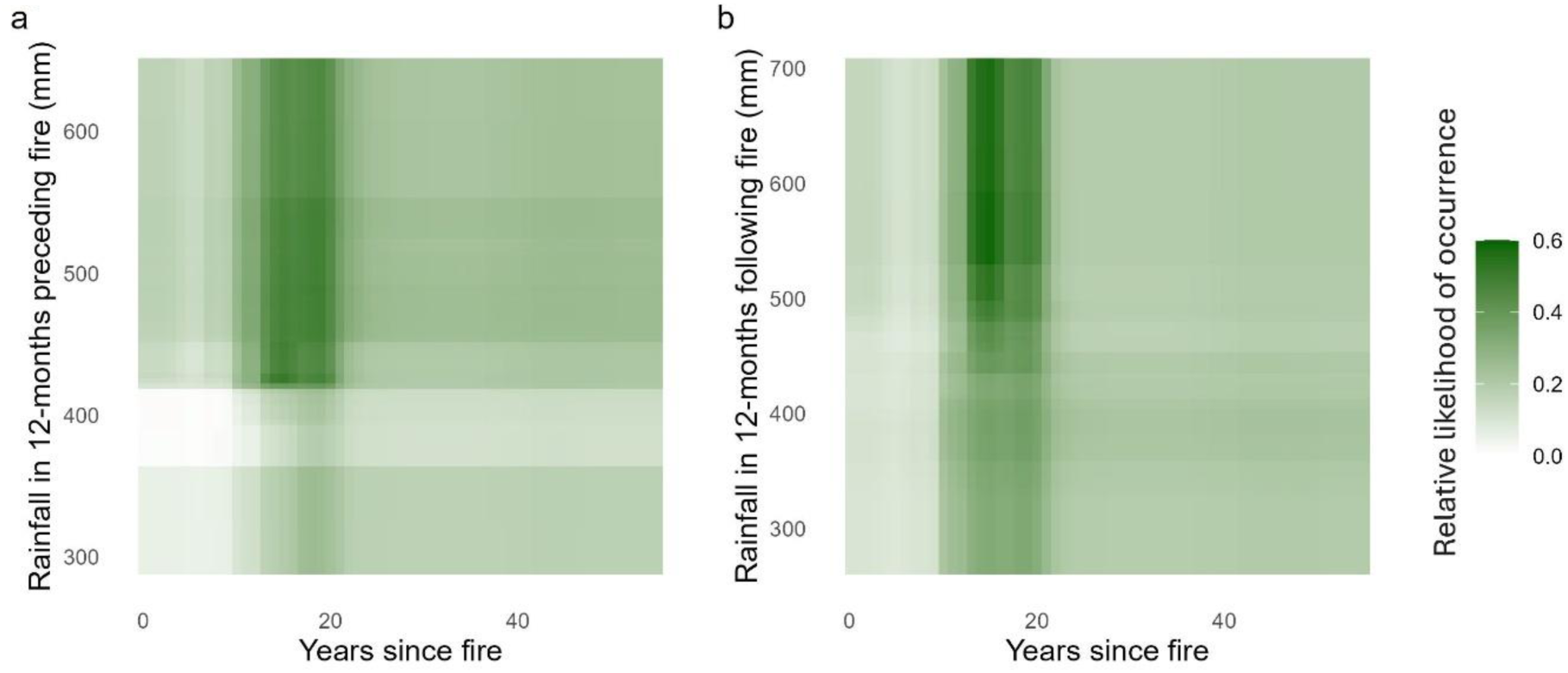
2-way partial dependence plots (PD) showing predicted relative likelihood of mallee emu-wren occurrence as a response to a) years since fire vs rainfall in the 12-months preceding fire and b) years since fire vs rainfall in the 12-months following fire.

The effect of rainfall in the 12-months following fire was dependent on interactions with other variables, particularly mean annual rainfall (Figures 2, 3 and Supplementary Figure S1). Greater rainfall in the 12-months following fire led to increased relative likelihood of mallee emu-wren occurrence in the drier parts of the landscape, yet the opposite was true for more mesic areas in the southern Ngarkat. In addition to these variables associated with fire regime, localised elevation, mean annual rainfall, aspect, and thorium content were ranked important predictors of mallee emu-wren occurrence according to either mean decrease in accuracy or mean decrease in Gini impurity (Figures 2 and 4).

**Figure 4.**
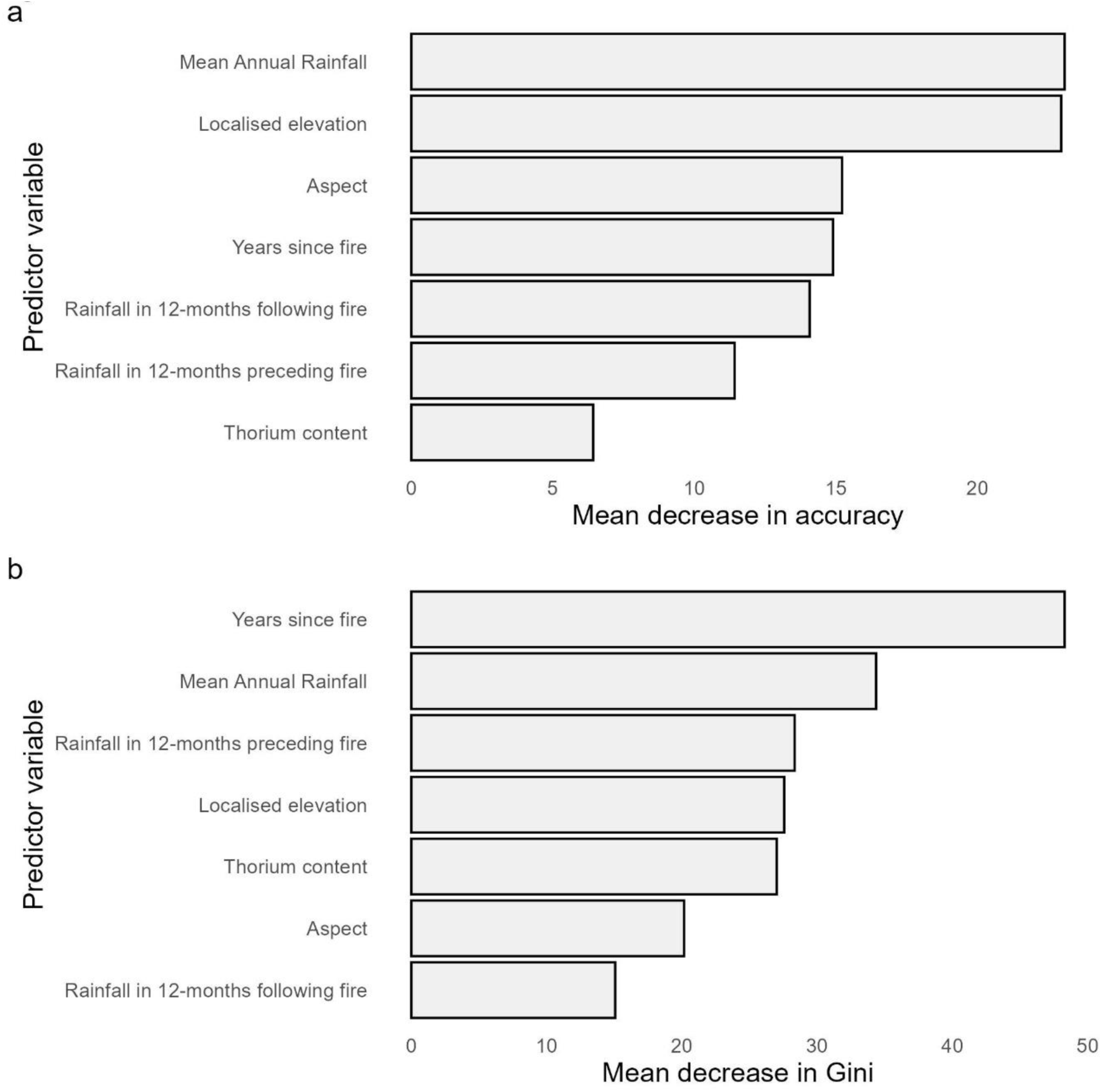
Order of importance of predictor variables included in a random forest classifier. Variable importance is ordered by a) mean decrease in accuracy (Does random variation of each variable reduce accuracy of the model?), and b) mean decrease in Gini impurity (how well does each variable contribute to reducing class mixing across all decision trees in a random forest?). For both ‘mean decrease in accuracy’ and ‘mean decrease in Gini impurity’, higher values indicate that a given variable has greater importance.

In December 2025, approximately 6,848 ha, or 2.5% of Ngarkat, was predicted to have a combination of environmental characteristics and fire-history favourable for mallee emu-wrens (predicted relative likelihood of occurrence > 0.5, Figure 5). Much of this potential habitat was concentrated in the south of the study region and was most recently burnt in 2014 (by wildfire). In the absence of future fire, 6,728 ha and 6,678 ha of Ngarkat are predicted to be favourable for mallee emu-wrens in 2035 and 2045 respectively (Figure 5).

**Figure 5.**
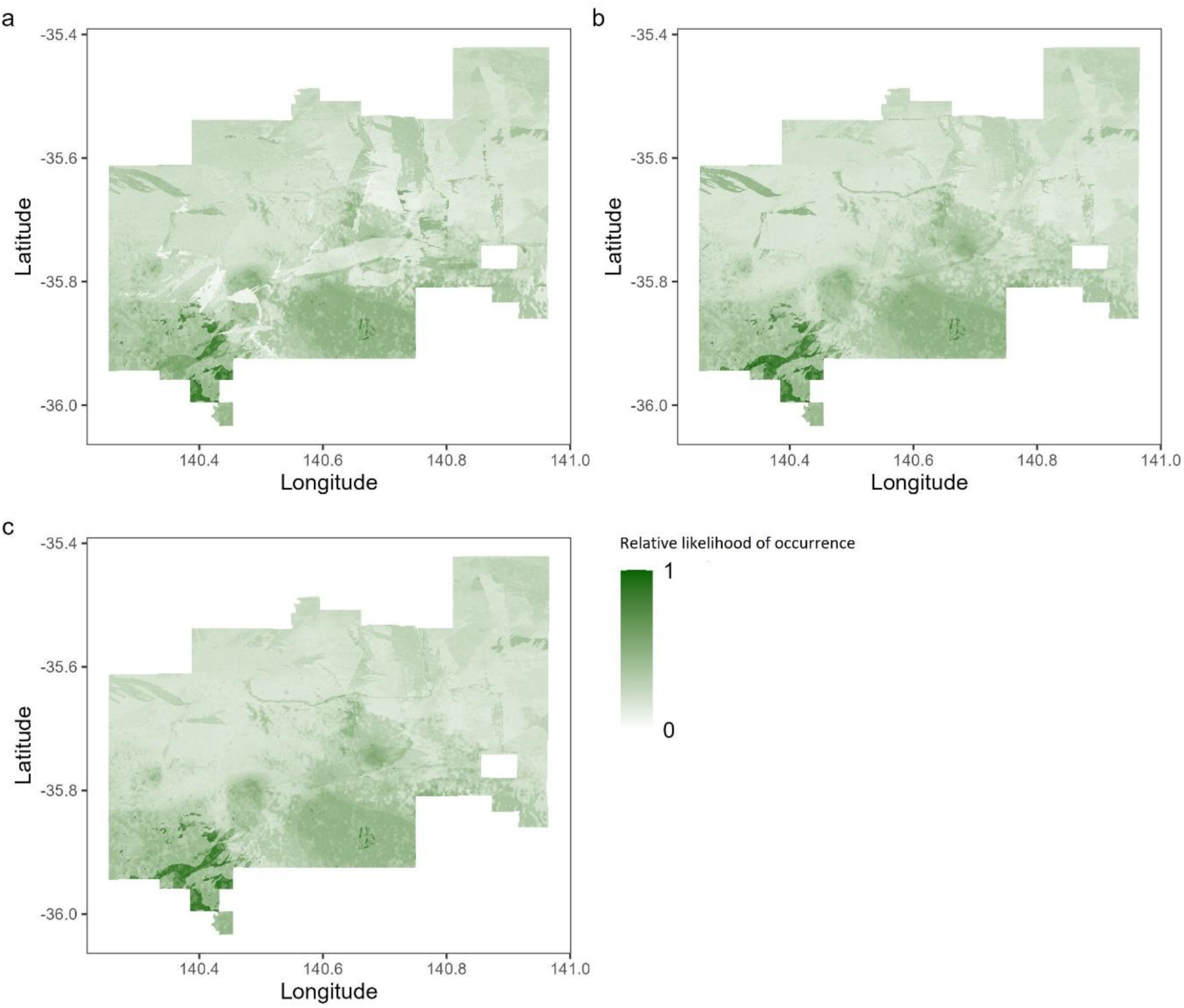
Predicted relative likelihood of mallee emu-wren occurrence across Ngarkat Conservation Park in a) December 2025, b) December 2035 and c) December 2045. This model used historical presence records collected prior to the local extinction of the species from this area. Here, relative likelihood of occurrence is considered a proxy for potential extant habitat.

## Discussion

Our results show that abiotic conditions around the time of a fire are likely to confer a long-term influence on post-fire occurrence for a rare, fire-sensitive species. Years since fire is an important predictor of mallee emu-wren relative likelihood of occurrence in heathland systems and mallee emu-wrens favour mid-successional vegetation (10 – 20 years post-fire). Fire is essential for maintaining mallee emu-wren habitat in this landscape and cumulative rainfall in the 12-month period before and after fire was associated with increased relative likelihood of occurrence for the mallee emu-wren ten 10 – 20 years post-fire. This result highlights conservation management challenges, and opportunities, that until now had not been identified. By managing fire such that most vegetation is burnt after favourable rainfall conditions (e.g., by prioritising management burning after periods of elevated rainfall and prioritising wildfire suppression efforts after drought in areas of ecological importance), managers have capacity to increase habitat quality for the mallee emu-wren over decades. In addition, such interventions would also benefit other taxa which share key habitat associations with the mallee emu-wren in Mediterranean shrubland ecosystems (Verdon et al. 2020, Verdon et al. 2025). Our results suggest that more temperate areas (i.e., southern Ngarkat) support mallee emu-wrens for longer and conservation interventions focused in such areas should be prioritised. Furthermore, our results identify a large area of south-central Ngarkat that is predicted to support mallee emu-wren habitat in 2025, and which may continue to provide habitat until at least 2045, if managed effectively. Identifying such locations is an important step toward the future reintroduction of this species to Ngarkat.

Conservation managers should first seek to protect areas of suitable habitat. However, in those areas that do not currently support taxa of conservation concern, or in which extant habitat is beginning to decline in quality, prioritising management burns after periods of above-average rainfall may facilitate regeneration of higher quality habitat for the mallee emu-wren, potentially expanding the proportion of heathland habitat suitable for the species. In heathland vegetation, the mallee emu-wren favours mid-successional vegetation, and some level of fire in the landscape will be required to ensure its ongoing persistence. In recent decades, most vegetation in this landscape has burnt after periods of below average rainfall and careful management would be required to reverse this trend (e.g., by increasing fire suppression during drought and burning during wetter periods). Although there may be challenges associated with burning during wetter periods (e.g., reduced capacity of fuel to sustain fire and achieve sufficient spread), achieving this goal in areas with other traits associated with the presence of mallee emu-wren habitat (e.g., areas characterised by heathland vegetation, low elevation, and more temperate climate) has potential to improve conservation outcomes in this landscape.

The potential benefits of a policy that prioritises burning during wetter periods are not limited to the mallee emu-wren. For example, vegetation which supports a high density of mallee emu-wrens typically overlaps with hotspots of other bird species of conservation concern, including the endangered Murray Mallee striated grasswren *A. striatus howei* and endangered red-lored whistler *Pachycephala rufogularis* (Verdon and Clarke 2022, Verdon et al. 2025). All three of these species share an association with *T. scariosa* which is considered a foundation species due to the resources and complex habitat it provides for a wide range of fauna (Moseby et al. 2016, Verdon et al. 2020, Bell et al. 2022). Relative to other ecosystems, Australian heathlands are typically able to carry fire within a relatively short time following substantial rain (Keith et al. 2002). This trait has been attributed to greater exposure to wind and sun compared with areas protected by an extensive canopy. If heathy vegetation has experienced dieback, the retention of dead limbs may also contribute to rapid drying and high flammability, even after periods of high rainfall (Keith et al. 2002). Identifying the optimum timing of management burns, such that vegetation is sufficiently dry to carry fire but while still enhancing ecological outcomes, will require further study.

Cumulative rainfall in the 12-months after fire also improves long-term habitat quality for mallee emu-wrens in heathland vegetation. The strength of this effect was dependent on mean annual rainfall, with post-fire rainfall conferring the greatest benefit in the driest parts of the landscape. *Triodia scariosa* is a core habitat feature for the mallee emu-wren with width, volume and percentage cover of *T. scariosa* shown to be important predictors of mallee emu-wren occurrence in Triodia mallee vegetation (Verdon et al. 2020). The positive effect of rainfall on relative likelihood of mallee emu-wren occurrence, highlighted in this study, may be mediated by the presence and condition of *T. scariosa*. *Triodia scariosa* relies on fire for recruitment (Rice and Westoby 1999). Recruitment can occur through either resprouting or reseeding after fire and, in either case, is positively affected by post-fire rainfall (Noble and Vines 1993, Giljohann et al. 2017). Much of the research investigating *in-situ* recruitment of *T. scariosa* (Cohn and Bradstock 2000, Giljohann et al. 2017, Bell et al. 2022) has occurred in regions that receive considerably lower median annual rainfall (<350 mm) than this study (327 – 451 mm). Mean annual rainfall has been shown to positively influence *T. scariosa* height and percentage cover (Kenny et al. 2018). It’s possible that in the southernmost part of Ngarkat, where we found post-fire rainfall had the least effect on relative likelihood of mallee emu-wren occurrence, high median annual rainfall diluted any positive association between post-fire cumulative rainfall and relative likelihood of mallee emu-wren occurrence.

This study highlights that a considerable area within Ngarkat is likely to support mallee emu-wren habitat. This species has limited dispersal capability and is unlikely to recolonise burnt areas without contiguous habitat connected to an extant population (Brown et al. 2013). A 2018 translocation of mallee emu-wrens to an area of Ngarkat that previously supported the species did not result in the establishment of a population, likely due to a period of drought following releases and high levels of dispersal (Mitchell et al. 2021). However, rainfall at the release area in the 12-months prior to most recent fire was below the 420 mm threshold identified in this study, suggesting that the quality of habitat may have been relatively poor. This study predicts that Ngarkat contains sufficient habitat to support a reintroduction of mallee emu-wrens and that core habitat in the southern part of Ngarkat should continue to support mallee emu-wrens until at least 2045. It also provides some insight into the management interventions that may facilitate the maintenance of that habitat under future climate change (i.e., ensuring management burns of senescing habitat occur after periods of elevated rainfall). The alignment of fire and favourable weather conditions to bolster the quality of regenerating habitat may be necessary to ensure the long-term persistence of any translocated population in Ngarkat.

### Potential mechanisms driving the association between rainfall at the time of fire and relative likelihood mallee emu-wren occurrence

Several potential pathways exist by which cumulative rainfall, either before or after fire, may influence habitat quality over the long-term (Figure 6). Elevated rainfall may increase the rate of fuel accumulation, increasing the probability and severity of fire (Clarke et al. 2021). This phenomenon is a typical feature of fire regimes in Triodia mallee vegetation, where continuous groundcover takes 10-20 years to develop (Bradstock and Cohn 2002, Clarke et al. 2021). In this system, elevated rainfall may promote the rapid growth of short-lived grasses, creating a continuous groundcover and temporarily increasing the likelihood and severity of fire. However, the heathland vegetation of Ngarkat has been more likely to burn after periods of below average rainfall, at least in recent decades. Elevated soil moisture and reduced desiccation associated with recent rainfall may reduce extent and severity of fire. Under such conditions, resprouting flora species typically have a higher rate of survival (Keith et al. 2002), which in turn may facilitate faster regeneration of habitat. Fire severity is likely a determinant of recruitment strategy for *T. scariosa* (Rice and Westoby 1999). High fire severity is more likely to kill individuals, resulting in proportionally greater recruitment from seed, while individuals are more likely to survive and recruit by resprouting under low fire severity.

**Figure 6.**
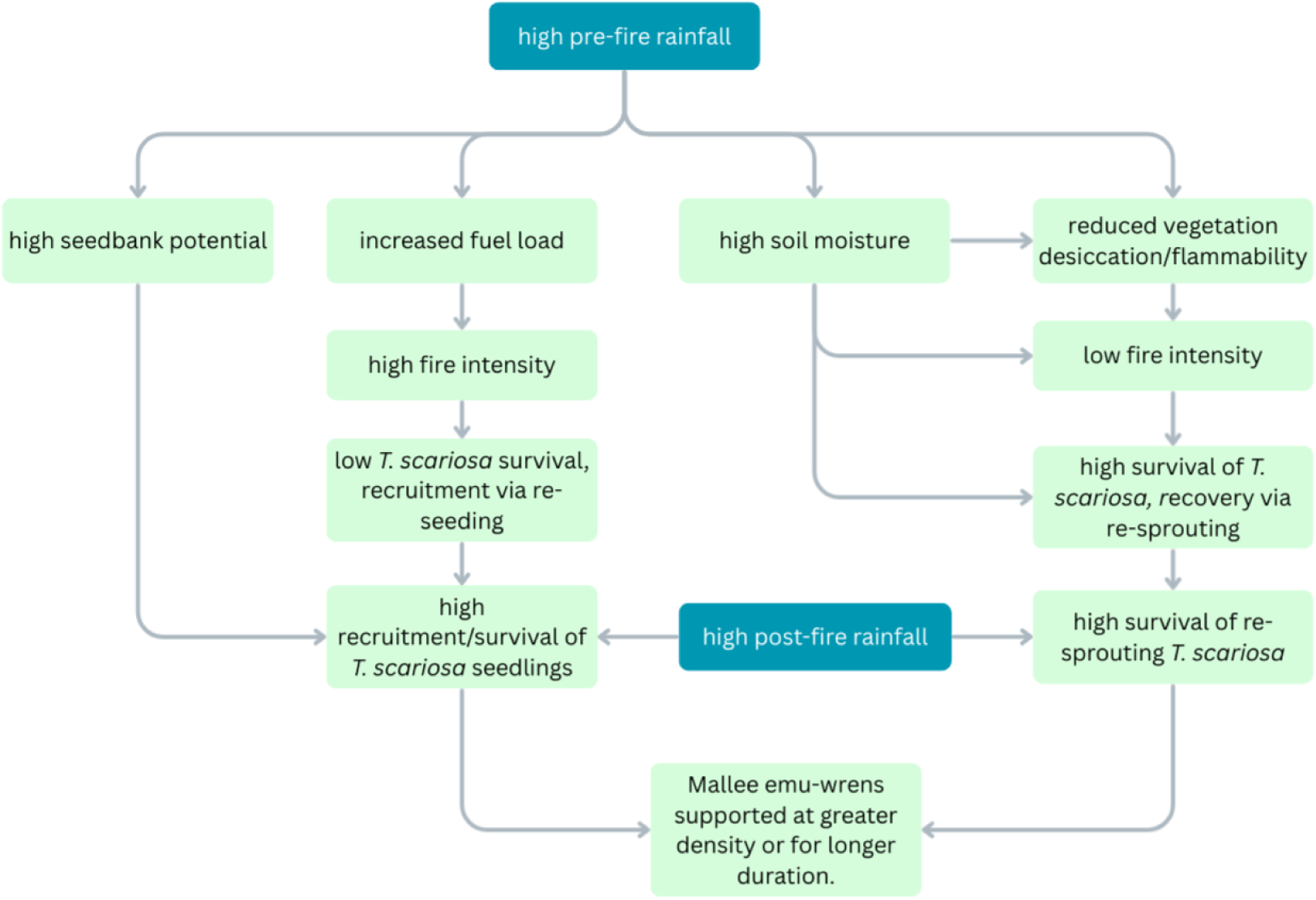
A conceptual diagram illustrating potential mechanisms linking rainfall before and after fire with long-term improved habitat quality for the mallee emu-wren. Note that the associations depicted in this figure represent hypotheses and are untested. This diagram does not include all potential drivers of mallee emu-wren habitat suitability (e.g., local climate, grazing pressure, interspecific competition or local geology).

Recent rainfall is likely to influence *T. scariosa* seedbank potential (Wright and Fensham 2018). In many *Triodia* species, including *T. scariosa*, masting events (mass synchronised seed crops) are triggered after periods of unusually high or sustained rainfall (Giljohann et al. 2017, Wright and Fensham 2018). Between such events, reproductive output is low (Wright and Fensham 2018). Seedbank dynamics are poorly understood in *T. scariosa* but in the congeneric soft spinifex *T. pungens*, a rainfall precipitated masting event resulted in a 200-fold increase in seedbank density (Wright and Fensham 2018). This spike was short-lived, lasting ∼24-months. *T. pungens* hummocks which were experimentally burned 6-months after this masting event showed significantly higher recruitment than control areas that were also burnt but at which inflorescences were clipped prior to seeding (Wright and Fensham 2018). This experiment demonstrated the importance of the timing of fire in relation to rainfall-precipitated mass seeding events in *Triodia* spp. It is possible that a similar phenomenon is contributing to increased relative likelihood of mallee emu-wren occurrence in areas which received high cumulative rainfall in the 12-months before most recent fire in Ngarkat. It is also unclear whether similar processes are occurring for other habitat providing plants (e.g., *Leptospermum* spp. or *Allocasuarina* spp.). Additional research identifying the mechanism linking cumulative rainfall before and after fire with mallee emu-wren relative likelihood of occurrence will improve the capacity for Mediterranean shrubland systems to be managed to maximise habitat quality for species of conservation concern.

### Limitations

Given the scarcity of mallee emu-wrens in heathland habitat, historical records of the species from areas they have since been extirpated are of incredible value. However, the use of unstructured presence-only data brings certain limitations. Adequate sampling of the full range of environmental variables likely to influence occurrence of target species was not always achieved in this study. Quantifying the effects of rainfall at different temporal scales could not be achieved in this study. The scale of real-world habitat features in comparison to that of available spatial data may have been an additional source of bias. Within heathland vegetation, small pockets of dense *T. scariosa* often form at drainage points in the landscape (e.g., south-easterly facing slopes of dunes). Such patches may be small, but provide important ‘anchor points’ around which mallee emu-wren territories often form (Mitchell et al. 2021). A spatial resolution of 75 m x 75 m necessitated by available data may not always capture such fine-scale habitat features. Despite these challenges, the random forest model used in this study performed well with 83% of presence class predictions shown to be correct. The predicted fire response of mallee emu-wrens generated by this model followed a ‘bell’ distribution, similar to that recorded for other species in this landscape (Watson et al. 2012, Makdissi et al. 2024). Given the low density typical of mallee emu-wren populations, even at sites that contain high-quality habitat, perfect model performance is unlikely to be achieved, especially given the fact that areas of regenerating habitat require connectivity with extant populations to become occupied.

Vegetation in the south-western corner of Ngarkat changes from the heathlands and mallee shrublands and woodlands typical of Ngarkat to a more mesic dry-forest vegetation type wholly unsuitable for the mallee emu-wren. This change in vegetation type is poorly captured by the broad vegetation categories represented in the spatial data currently available. As such, some of the areas with high predicted relative likelihood of mallee emu-wren occurrence in the south-western corner of Ngarkat should be treated with caution. This example highlights the need for careful, well-informed and ecologically integrated application of SDMs in conservation management (Guillera-Arroita et al. 2015).

The mechanism underlying the association between rainfall at the time of fire and relative likelihood of mallee emu-wren occurrence remains unknown (Fig. 6). By framing future management around experimental trials of interventions designed to shed light on this mechanism (e.g., timing of fire, augmenting seedbanks after fire, and supplemental watering before or after fire), a greater understanding of fire-recovery dynamics in Mediterranean shrubland ecosystems may be achieved while improving future habitat for a suite of rare and threatened taxa, including the mallee emu-wren. An experimental study which included simulated rainfall through supplemental watering would increase the resolution of rainfall data at the site scale, allowing greater capacity to investigate the impact of variability in rainfall across the study area and to control for variability across long time periods. Such a study would provide the opportunity to incorporate other factors likely to influence the regeneration of key habitat taxa that were not able to be incorporated into this study (e.g., grazing pressure).

### Conclusions

Managing fire to promote populations of rare fire-sensitive species presents considerable challenges for conservation managers. This study informs effective fire management by identifying a previously unrecognised association between the long-term quality of regenerating habitat and distinct abiotic events, namely cumulative rainfall in the 12-months before and after the most recent fire. The impact of rainfall before and after the most recent fire on long-term habitat quality presents both challenges and opportunities for conservation management of the mallee emu-wren. Although operational factors may present challenges (difficulty burning after wetter periods), fire managers have capacity to increase the future extent of mallee emu-wren habitat in Ngarkat by burning during/after wetter periods and suppressing fire during dry periods.

More broadly, the abiotic conditions around the time of a fire represent an important and under-appreciated driver of post-fire recovery over the long-term, potentially acting in many ecosystems globally. Our capacity to deliver effective fire management for habitat regeneration requires a better understanding of how abiotic conditions around the time of a fire influence post-fire recovery in a range of fire-prone ecosystems.

The authors have no conflict of interest to declare.

## Notes

### Competing Interest Statement

The authors have declared no competing interest.

