## Supplementary Material for "Rainfall immediately before and after fire promotes long-term occurrence of a rare, fire-sensitive passerine"

### Variability in model performance between Ngarkat Conservation Park and the adjacent Big Desert complex

There was a western bias in species records available for this study. Only 25 presence records, and 273 background points were available from the Big Desert complex (Big Desert) in the state of Victoria, connected to and located east of the Ngarkat Conservation Park (Ngarkat) study area (South Australia). As such, we expected that predictive performance of any model trained on this dataset was likely to be better in Ngarkat than Big Desert. To test this, our random forest classifier was trained using only data from Ngarkat. We assessed model performance using out of bag (OOB) error rate. We then predicted probability of mallee emu-wren presence at each of 298 Big Desert data points and compared predictions against the true state to calculate error rate for both positive (presence) and negative (background) classes. A second test was used to further validate model performance in both Ngarkat and Big Desert. During a large-scale survey in 2022 (Verdon et at. 2025), experienced ecologists assessed mallee emu-wren habitat suitability at 229 25-ha sites across the study region. Each site consisted of six equal-sized cells (4.2-ha). For each cell, an experienced ecologist assigned one of three qualitative categories regarding the presence of mallee emu-wren habitat: no habitat, marginal habitat or good habitat. Using this dataset, we assigned a simple habitat score to each site. If any of one of the six cells was scored as 'good habitat', we considered mallee emu-wren habitat to be suitable at the site scale. We then used the random forest classifier to predict the probability of mallee emu-wren presence at the centre point of each plot and compared mean probability of presence (based on the RF classifier) between sites judged to either support or not support good mallee emu-wren habitat. We did this separately for both Ngarkat and Big Desert and used two-sample t-tests to check for a significant difference between sites with no habitat and good habitat.

The RF classifier developed in this study had greater predictive performance in Ngarkat compared with the Big Desert. In Ngarkat, the random forest model predicted mallee emu-wren presence with an error rate of 0.20 and absence with an error rate of 0.10. In Big Desert, the same model predicted mallee emu-wren presence with an error rate of 1 (i.e., no presences were successfully detected) and absences with an error rate of 0.01. Our second validation test, comparing predicted probability of mallee emu-wren presence between sites where extant habitat was considered by experts to be either suitable or not suitable, yielded similar results. In Ngarkat, predicted probability of mallee emu-wren presence was significantly higher ($T_{44} = 3.44$, $p = 0.001$) at plots judged to contain suitable mallee emu-wren habitat when compared with plots that did not contain suitable habitat (Table S1). By contrast, no significant difference in predicted probability of mallee emu-wren presence was detected between sites with habitat judged as suitable or not suitable in Big Desert (Table 2; $T_{181} = 1.09$, $p = 0.278$). This result suggests that the environmental conditions underpinning suitable mallee emu-wren habitat likely differ between the contiguous Ngarkat and Big Desert, despite sharing broadly similar vegetation. Based on these results, we limited the scope of this study to the 271,000 ha Ngarkat Conservation Park.

Table S1. Probability of mallee emu-wren occurrence, predicted using a random forest (RF) classifier, at habitat plots qualitatively categorised as either suitable or not suitable. Performance of the RF classifier is compared Ngarkat and Big Desert.

| Qualitative habitat assessment | Mean predicted probability of mallee emu-wren presence ± se | |
|---|---|---|
| | Big Desert | Ngarkat |
| Not suitable for mallee emu-wren | 0.31 ± 0.01 | 0.40 ± 0.02 |
| Suitable for mallee emu-wren | 0.33 ± 0.02 | 0.60 0.09 |

Figure S1. 2-way partial dependence plot depicting a) interaction between annual rainfall (y axis) and cumulative rainfall in the 12-month period preceeding fire (x axis), with predicted probability of mallee emu-wren presence as response (colour gradient) and b) interaction between annual rainfall (y axis) and cumulative rainfall in the 12-month period following fire (x axis), with predicted probability of mallee emu-wren presence as response (colour gradient).

A)

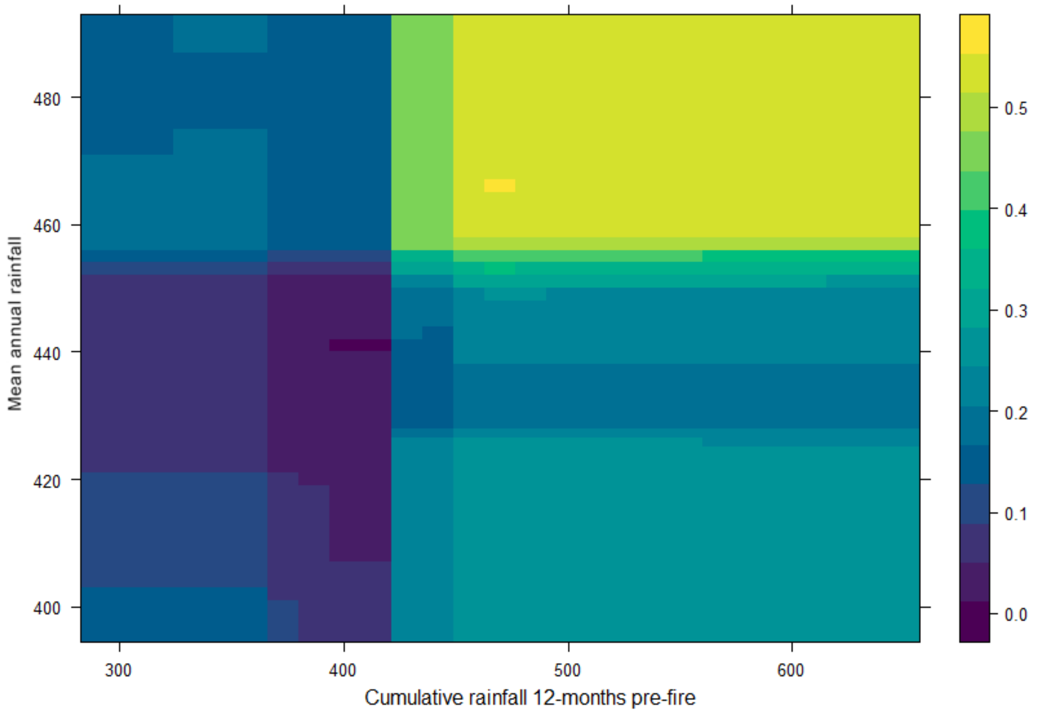

B)

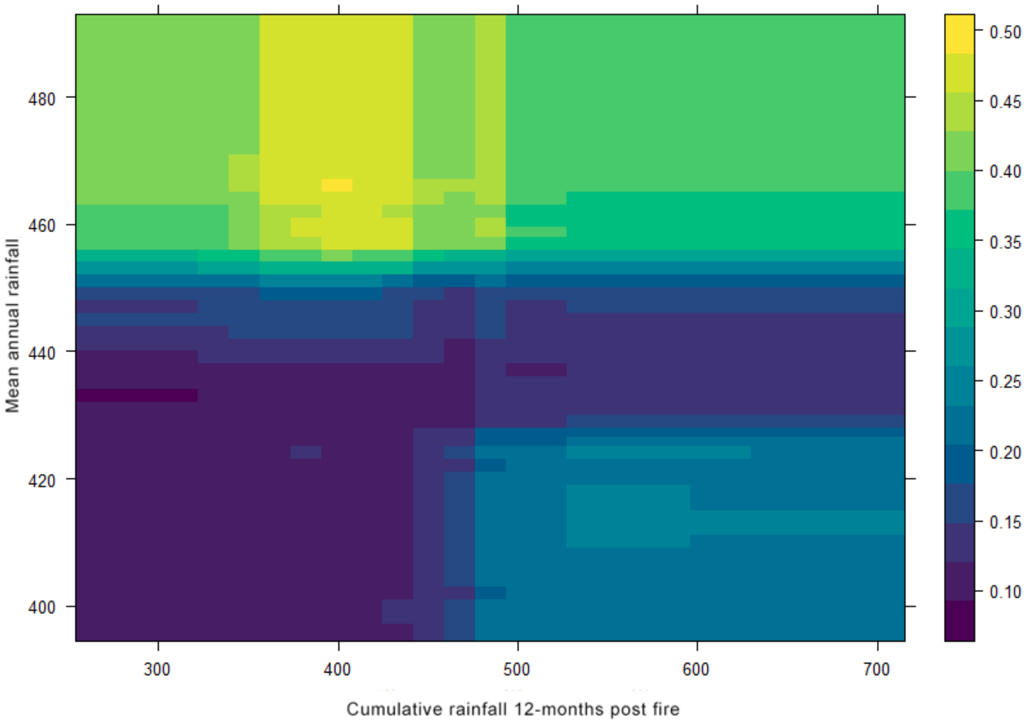

Mean annual rainfall
480
460
440
420
400
300
400
500
600
700
Cumulative rainfall 12-months post fire
0.50
0.45
0.40
0.35
0.30
0.25
0.20
0.15
0.10
